# Phage display-mediated immuno-PCR to detect low-abundance secreted proteins in *Drosophila*

**DOI:** 10.1101/2025.10.31.685841

**Authors:** Myeonghoon Han, Baolong Xia, Ah-Ram Kim, Elizabeth Filine, Emily Stoneburner, Ting Miao, Ying Liu, Jonathan Zirin, Norbert Perrimon

## Abstract

Circulating hormones, that mediate communications across organs to maintain physiological balance, are commonly detected and quantified using enzyme-linked immunosorbent assays (ELISAs). However, while ELISA is well-suited for organisms where sample blood can be readily obtained, its application is considerably more challenging in smaller organisms, particularly *Drosophila*, which has gained widespread use in recent years for physiological studies. Here, we present sensitive phage display-mediated immuno-PCR (PD-iPCR) to detect *Drosophila* hemolymph proteins via two approaches: 1) by identifying high-affinity nanobodies through phage-display library screening and subsequent affinity maturation and 2) by generating a knock-in fly line producing secreted proteins tagged with tandem NanoTags composed of VHH05 and 127D01. Using these approaches, we successfully established PD-iPCR to detect insulin-binding ImpL2 protein in fly hemolymph. Notably, the tandem NanoTag-based sandwich PD-iPCR enabled highly sensitive detection of tagged antigens, allowing us to quantify elevated ImpL2 levels in the hemolymph of starved flies and those bearing *Yki*-induced gut tumors. Collectively, our results demonstrate that PD-iPCR enables detection of endogenous, low-abundance circulating hormones in *Drosophila*, providing a powerful tool for studying interorgan communication.

**Significance statement:** Hormones and other secreted factors orchestrate organism-wide physiology, yet their routine quantification in *Drosophila* has been limited by the limited volume of hemolymph available for assays like enzyme-linked immunosorbent assay (ELISA). Here, we present phage display-mediated immuno-PCR (PD-iPCR) as a novel sensitive platform for quantifying secreted proteins in flies *in vivo*. Using ImpL2 as an example, we successfully detected nanomolar level of circulating ImpL2 and monitored its physiological changes during starvation and tumorigenesis using PD-iPCR. This approach can be readily expanded to multiplexed quantification of secreted proteins *in vivo* by leveraging the multiple available nanobodies and the vast collection of epitope-tagged *Drosophila* lines. This work opens the door to systematic endocrine phenotyping across developmental stages and diverse physiological conditions.

## Introduction

Circulating factors in the blood, such as hormones, growth factors, and cytokines, play essential roles in coordinating communication between organs and maintaining systemic physiological balance. These signaling molecules respond dynamically to internal and external cues, regulating processes such as growth, metabolism, reproduction, and stress responses. For example, insulin, a key hormone that regulates blood glucose levels, is tightly regulated in response to nutrient intake (1). After meals, insulin secretion increases to promote glucose uptake and storage, while fasting reduces insulin levels. Measuring circulating insulin is standard practice in both clinical and research settings to assess metabolic health, diagnose disorders such as diabetes, and monitor therapeutic responses.

Enzyme-linked immunosorbent assays (ELISAs) are widely used to detect and quantify circulating hormones due to their specificity, scalability, and relative ease of use. While ELISA is well-suited for organisms such as humans or mice, where sample blood volume and target protein can be readily obtained from a single individual, its application is considerably more challenging in smaller organisms. One such case is *Drosophila melanogaster*, a powerful genetic model that has gained widespread use in recent years for studying metabolism, physiology, and inter-organ communication (2, 3). Major technical limitations in flies include the extremely small volume of hemolymph (the fly equivalent of blood), typically less than 100 nL per individual (4), and the low abundance of circulating proteins (5), both of which present substantial obstacles for conventional ELISA.

To date, only very few studies have quantified secreted proteins in *Drosophila*, including ELISA-based detection of *Drosophila* insulin-like peptide 2 (Dilp2) (6) and Adipokinetic hormone (Akh), a glucagon-like hormone (7). Therefore, developing more sensitive and broadly applicable methods for quantifying circulating proteins in *Drosophila* remains a pressing need in the field. Such tools would enable precise monitoring of hormonal and metabolic signals at physiological concentrations in individual flies or small cohorts, expanding our ability to explore systemic regulation in this important model organism.

Here, we present a phage display-mediated immuno-PCR method for the detection of circulating hormones in *Drosophila*. Using *Drosophila* ImpL2 as a representative example, we quantified ImpL2 in hemolymph via two approaches: 1) identification of high-affinity nanobodies that selectively bind endogenous ImpL2, and 2) generation of a transgenic fly line in which tandem NanoTags are knocked into the endogenous *ImpL2* genomic locus. Using these tools, we successfully detected endogenous ImpL2 protein in the hemolymph and observed changes in circulating ImpL2 levels under various physiological conditions. Together, these approaches offer robust and sensitive strategies for measuring low-abundance fly hormones.

## Results

### ImpL2 nanobody screening using phage-displayed nanobody library

ImpL2 binds to Dilp2 and Dilp5 thereby systemically suppressing insulin signaling (8, 9). Several groups have generated antibodies to capture the endogenous ImpL2 protein, by which several key biological functions of ImpL2 have been studied (10–15). However, the resources to measure endogenous ImpL2 remain limited, and incapable of estimating ImpL2 molarity. To address this issue, we decided to generate nanobodies against ImpL2 using a phage-displayed nanobody library and screening platform that we recently established (16). To do this, we expressed a human IgG Fc domain-fused ImpL2 (ImpL2-hIgG) in *Drosophila* S2 cells, ensuring that it was secreted via native protein maturation and secretion pathways. ImpL2-hIgG was then purified from conditioned medium using Protein A resin. Nanobody screening was carried out through three rounds of selection (Figure 1A; see Materials and Methods). Following three rounds of selection, the outcomes were assessed by ELISA using polyclonal phages collected after each round. As expected, the ELISA signal increased after three rounds of selection, indicating progressive enrichment of phages displaying nanobodies with affinity for the antigen in the polyclonal population (Figure 1B). Next, we identified positive hits from the nanobody screens by ELISA using monoclonal phages isolated after the third round of selection, selecting phages that showed strong signals against ImpL2-hIgG but not the control protein, mCherry-hIgG (Figure 1C). Sanger sequencing of 23 positive clones revealed four unique nanobody sequences (Table 1).

**Figure 1.**
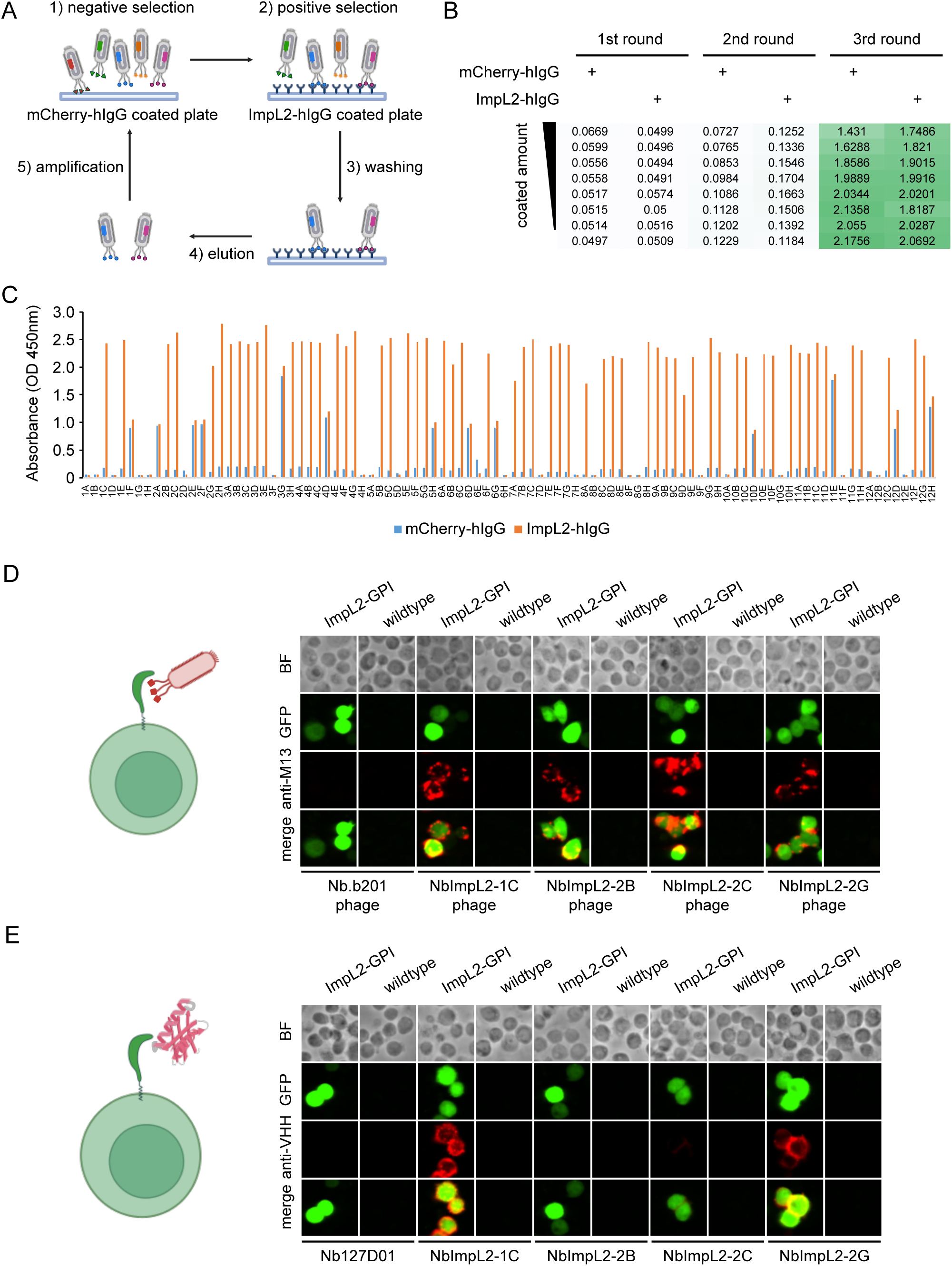
ImpL2 nanobody screening using phage-displayed nanobody library. (A) Schematic of ImpL2 nanobody screening. A Maxisorp plate coated with mCherry-hIgG was used for negative selection, and a coated plate with ImpL2-hIgG was used for positive selection. Three iterative rounds of selection were conducted. (B) ELISA results using polyclonal phages from three rounds of selection. The coating amount of mCherry-hIgG and ImpL2-hIgG decreased across the rows. The intensity of the green color indicates the strength of the ELISA signal. (C) ELISA results of monoclonal phages isolated after the third round of selection. 96 individual clones were tested against the control protein (mCherry-hIgG) and the target protein (ImpL2-hIgG). (D) Immunostaining of cells expressing membrane-tethered ImpL2 using nanobody-displaying phages. GFP signals mark cells expressing membrane-tethered ImpL2. Bound phages were detected using an antibody against the M13 major coat protein. Nb.b201-displaying phage was used as a negative control. BF, bright field. (E) Immunostaining of cells expressing membrane-tethered ImpL2 using recombinant nanobodies. GFP signals mark cells expressing membrane-tethered ImpL2. Bound nanobodies were detected using an antibody against alpaca VHH fragments. Recombinant Nb127D01 was used as a negative control. BF, bright field.

**Table 1.**
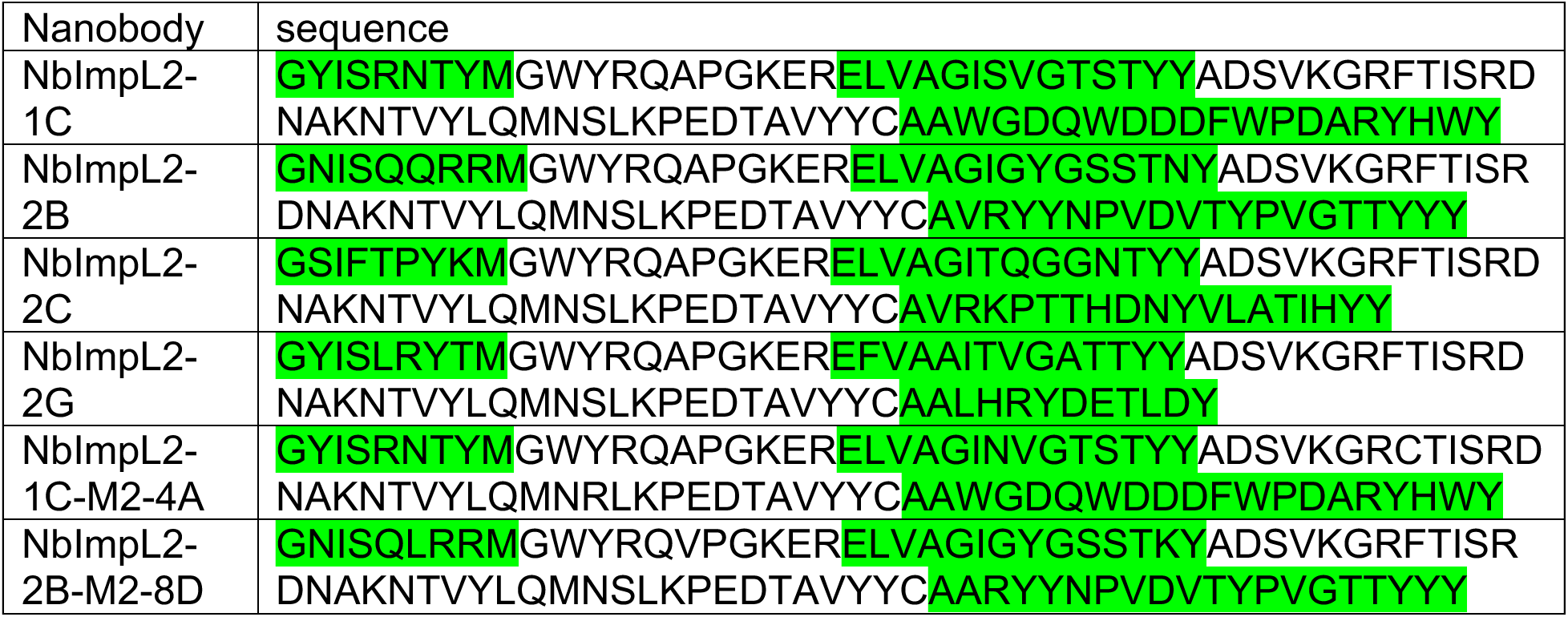
Protein sequences of ImpL2 nanobodies and their variants used in this study, with CDR regions highlighted in green.

To further assess whether the nanobody candidates specifically recognize ImpL2, we anchored ImpL2 to the cell surface using a glycosylphosphatidylinositol (GPI) tag and incubated the cells with nanobody-displaying phages. Unlike an irrelevant nanobody Nb.b201, a nanobody against human serum albumin (17), all four ImpL2 nanobody-displaying phages bound robustly to ImpL2-tethered cells (Figure 1D). Additionally, purified recombinant nanobodies NbImpL2-1C and NbImpL2-2G showed clear binding to ImpL2-expressing cells (Figure 1E), further confirming their specificity for the target antigen. Taken together, these results demonstrate the successful identification of ImpL2 nanobodies using the phage-displayed nanobody library.

### ImpL2 nanobody affinity maturation by random mutagenesis

Since our initial assays used purified or overexpressed ImpL2, we next tested whether the nanobodies could recognize endogenous ImpL2 secreted by wild-type S2R+ cells using immunoprecipitation. However, only minimal amounts of endogenous ImpL2 were immunoprecipitated by the nanobodies (Supplementary Figure 1A), suggesting that their affinities were relatively low.

To address the affinity issue, we performed affinity maturation on all four ImpL2 nanobodies using error-prone PCR to introduce random mutations in both the complementarity-determining regions (CDRs) and framework regions. For each candidate, we generated a mutant library of approximately one million variants for additional rounds of screening. After two rounds of mutagenesis (see Materials and Methods), we observed several monoclonal phages showing higher ELISA signals than their original nanobody displaying phages (Figure 2A and Supplementary Figure 1B-1G), suggesting an increase in affinity.

**Figure 2.**
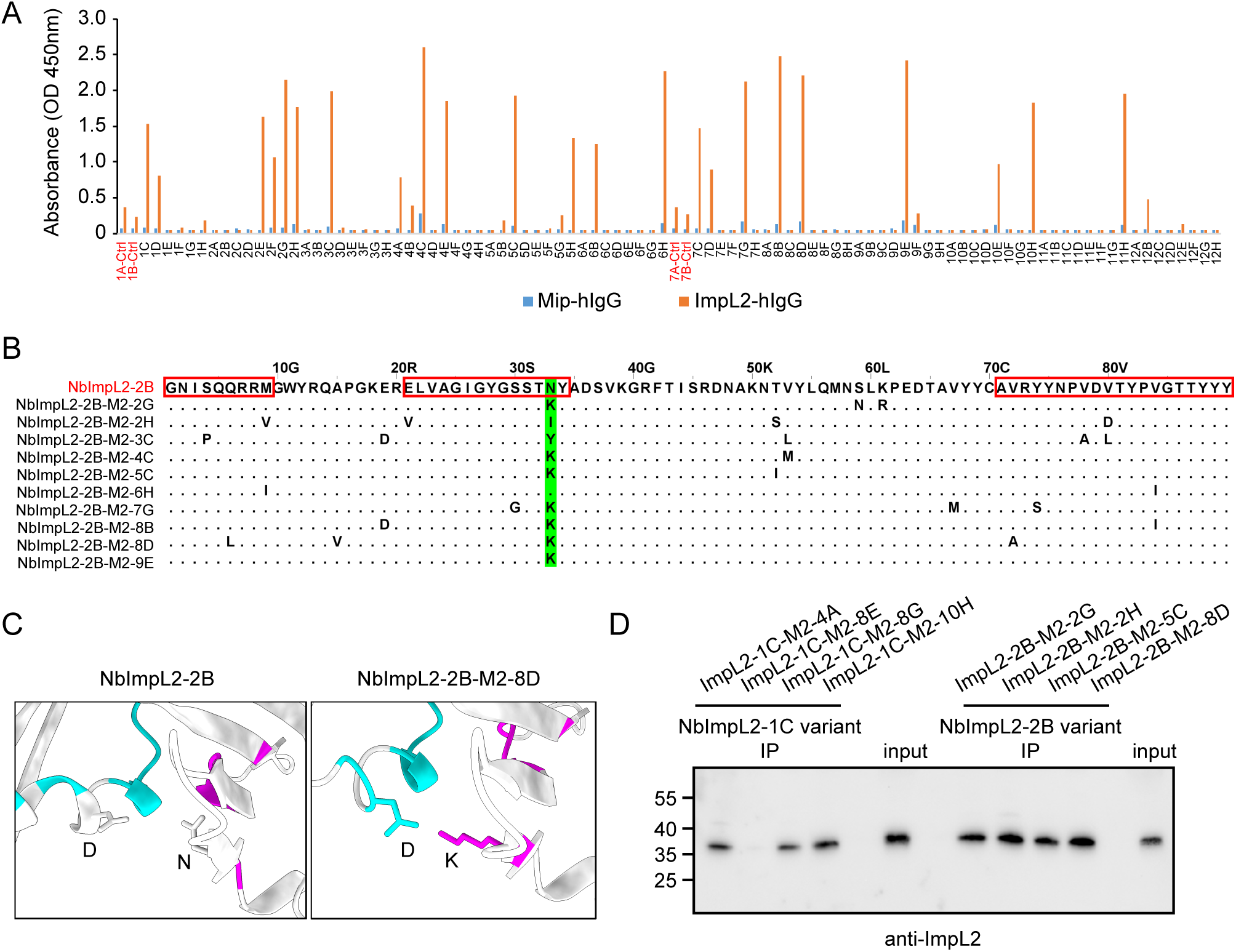
ImpL2 nanobody affinity maturation by random mutagenesis. (A) ELISA results of monoclonal phages isolated after two rounds of mutagenesis and selection of NbImpL2-2B. 96 individual clones were tested against the control protein (Mip-hIgG) and the target protein (ImpL2-hIgG). Clone 1A, 1B, 7A, 7B are phages displaying the original NbImpL2-2B nanobody. (B) Sequence alignment of NbImpL2-2B variants showing improved ELISA signals in (A). Mutations in the variants are indicated. The CDR regions are outlined with red rectangles, and the amino acids corresponding to asparagine residue in CDR2 are highlighted in green. (C) AlphaFold-Multimer prediction of the interaction between ImpL2 and either NbImpL2-2B or its variant NbImpL2-2B-M2-8D. Local interaction residues in contact interface (predicted aligned error ≤ 12 Å and Cβ–Cβ contact ≤ 8 Å) are highlighted in cyan for ImpL2 and magenta for the nanobodies. (D) Immunoprecipitation of ImpL2 from conditioned medium using NbImpL2-1C and NbImpL2-2B variants. Four different variants for each nanobody were tested. The ImpL2 was detected using endogenous ImpL2 antibody.

Notably, for the affinity maturation of NbImpL2-2B, among the 10 variants with improved ELISA signal, 9 carried mutations at an asparagine residue in CDR2—7 of which were substitutions to lysine (N-to-K) (Figure 2B). This strongly suggests that the asparagine in CDR2 limits the original nanobody’s affinity, and the N-to-K mutation enhances its binding to ImpL2.

To investigate the mechanism underlying the enhanced affinity conferred by the N-to-K mutation in NbImpL2-2B, we modeled the ImpL2-nanobody complex using AlphaFold-Multimer-based structural predictions (18). The predicted 3D structure supported our findings, revealing that the positively charged lysine side chain in CDR2 of NbImpL2-2B-M2-8D (a variant of NbImpL2-2B, NbImpL2-2B denotes the parental nanobody, M2 indicates two rounds of mutagenesis, and 8D refers to the specific clone selected after mutagenesis) forms favorable interactions with a negatively charged aspartic acid residue in ImpL2 (Figure 2C). This interaction was absent in the complex with the original NbImpL2-2B, explaining the observed increase in binding affinity. Importantly, after affinity maturation, NbImpL2-1C and NbImpL2-2B variants—but neither NbImpL2-2C nor NbImpL2-2G variants—efficiently immunoprecipitated endogenous ImpL2 from conditioned medium of S2R+ cells (Figure 2D). In summary, through nanobody screening and affinity maturation, we successfully identified high-affinity nanobodies against ImpL2.

### ImpL2 quantification using phage-display mediated immuno-PCR

Immuno-PCR combines the specificity of antibody-based detection with the high sensitivity of PCR amplification by using DNA-conjugated antibodies to recognize target molecules and amplify detection signals (19). Compared to conventional ELISA using horseradish peroxidase (HRP)-conjugated antibodies, immuno-PCR offers significantly greater sensitivity and a broader linear dynamic range (19). Building on this concept, previous study developed phage display-mediated immuno-PCR (PD-iPCR), which leverages phage-displayed single-chain variable fragments (scFvs) as detection antibodies, and utilizes the phage’s single-stranded DNA genome as the PCR template (20). In our system, we adapted this approach using nanobody-displaying phages and observed approximately a 1,000-fold increase in sensitivity compared to ELISA (Figure 3A), underscoring the powerful signal amplification capability of PCR-based detection.

**Figure 3.**
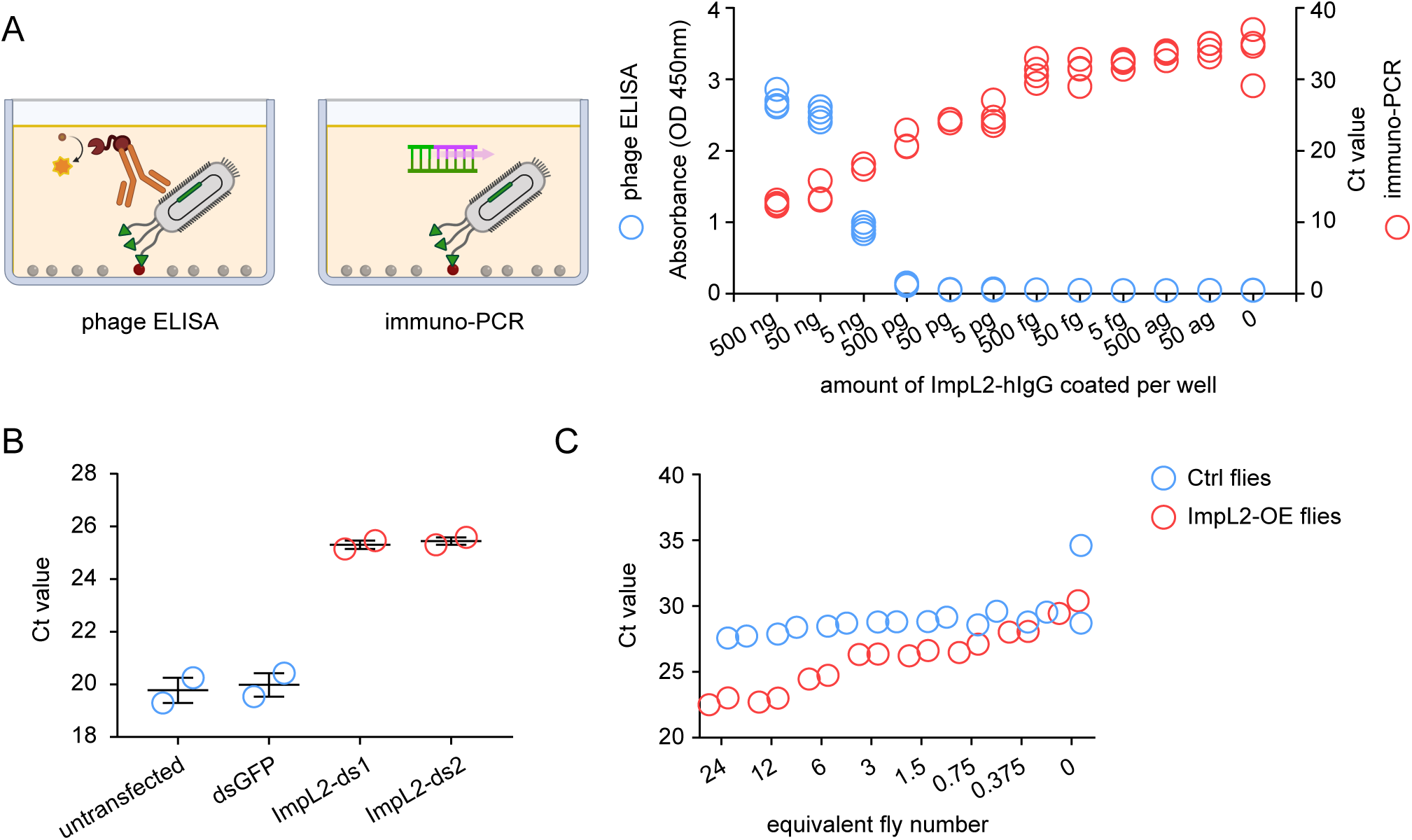
Immuno-PCR with ImpL2 nanobodies. (A) Comparison between phage ELISA and immuno-PCR. Purified ImpL2-hIgG was coated onto plates with a 10-fold serial dilution series. ImpL2 was detected using NbImpL2-2B-M2-8D-displaying phages by ELISA (blue circles) and immuno-PCR (red circles). Data represent four biological replicates. (B) Quantification of ImpL2 in conditioned medium from *Drosophila* S2R+ cells transfected with dsGFP or two distinct dsRNAs targeting *ImpL2*, measured by immuno-PCR. Data represent two biological replicates. Error bars, SEM. (C) Quantification of ImpL2 in the hemolymph of Ctrl flies (*Tub-Gal80^TS^/UAS-empty;mef2-Gal4/+*) and ImpL2-OE flies (*Tub-Gal80^TS^/+;mef2-Gal4/UAS-ImpL2*) using immuno-PCR. Flies were induced at 29 ℃ for 2 days. 2-fold serially diluted hemolymph samples were analyzed. Data represent two biological replicates.

Next, we tested whether sandwich PD-iPCR could be used to quantify ImpL2 levels in conditioned medium. We coated MaxiSorp plates with purified recombinant NbImpL2-2B variant, NbImpL2-2B-M2-8D as a capture nanobody and incubated the plates with conditioned medium from either control or *ImpL2* knockdown cells. To detect captured ImpL2, we used NbImpL2-1C variant, NbImpL2-1C-M2-4A-displaying phages and quantified the bound phage DNA via qPCR. Since higher levels of ImpL2 lead to increased phage binding, this results in lower cycle threshold (Ct) values in qPCR. As expected, conditioned medium from control cells showed markedly lower Ct values compared to *ImpL2* knockdown samples (Figure 3B), demonstrating that ImpL2 levels can be quantitatively measured using this method.

We then asked whether this approach could also be applied to quantify ImpL2 levels in fly hemolymph. After inducing *ImpL2* overexpression in fly muscle for two days, we collected hemolymph and measured ImpL2 levels using the same sandwich PD-iPCR setup. Hemolymph from ImpL2-overexpressing flies exhibited lower Ct values than that from control flies (Figure 3C). Additionally, serial dilution of hemolymph samples from *ImpL2*-overexpressing flies resulted in progressively increased Ct values (Figure 3C), confirming the sensitivity and dynamic range of the assay. Together, these results demonstrate that sandwich PD-iPCR enables sensitive and quantitative detection of ImpL2 in both cell culture supernatants and fly hemolymph.

ImpL2 induces systemic organ wasting in *yki^act^*-induced gut tumors in *Drosophila* (21). To determine whether the increase in ImpL2 could be detected in the hemolymph of flies bearing *yki^act^*-induced gut tumors, we collected hemolymph from both female and male flies after four and six days of *yki^act^* induction and measured ImpL2 levels using the sandwich PD-iPCR. However, no detectable increase in ImpL2 was observed in these samples (Supplementary Figure 2). We speculate that the sandwich PD-iPCR using ImpL2 nanobodies may not yet be sensitive enough to detect the elevated ImpL2 levels in the hemolymph of *Yki* flies.

### Detecting ImpL2 with tandem NanoTags using sandwich PD-iPCR

To overcome the limited sensitivity of our nanobody-based detection of endogenous ImpL2 in *Yki* flies (Supplementary Figure 2), we pursued an alternative strategy: tagging endogenous ImpL2 with detectable epitopes. Our recent work identified two NanoTag epitopes—VHH05 and 127D01—and their corresponding nanobodies (NbVHH05 and Nb127D01), which exhibit high specificity and functionality in *Drosophila in vivo* (22).

We first tested whether these NanoTags and nanobodies are compatible with ELISA. A synthetic peptide containing tandem NanoTags (referred to hereafter as tNTs; Figure 4A) was recognized by either NbVHH05 or Nb127D01 in direct ELISA, with detection limit at ∼100 fmol tNTs (Figure 4B). We then asked whether these tags could also be detected using sandwich ELISA, which typically offers greater sensitivity and specificity than direct ELISA. Using AlphaFold3 (23), we modeled the ternary complex of tNTs simultaneously bound by NbVHH05 and Nb127D01. The prediction suggested a stable complex formation (Figures 4C and 4D). To experimentally validate this, we transfected S2R+ cells with a plasmid encoding tNTs-fused GFP (GFP^tNTs^) to secrete into the culture medium. We then compared detection sensitivity between direct and sandwich ELISA for GFP^tNTs^. Sandwich ELISA showed superior sensitivity to detect GFP^tNTs^ than direct ELISA (Figure 4E). Notably, the signal was consistent regardless of the length of linker that combines two NanoTags (GFP^tNTs^_1×GSG, GFP^tNTs^_2×GSG, and GFP^tNTs^_3×GSG), suggesting that even a short three-amino-acid linker is sufficient to form a stable ternary complex. Swapping the capture and detection nanobodies between NbVHH05 and Nb127D01 also yielded comparable results (Figure 4E), confirming flexibility in assay configuration.

**Figure 4.**
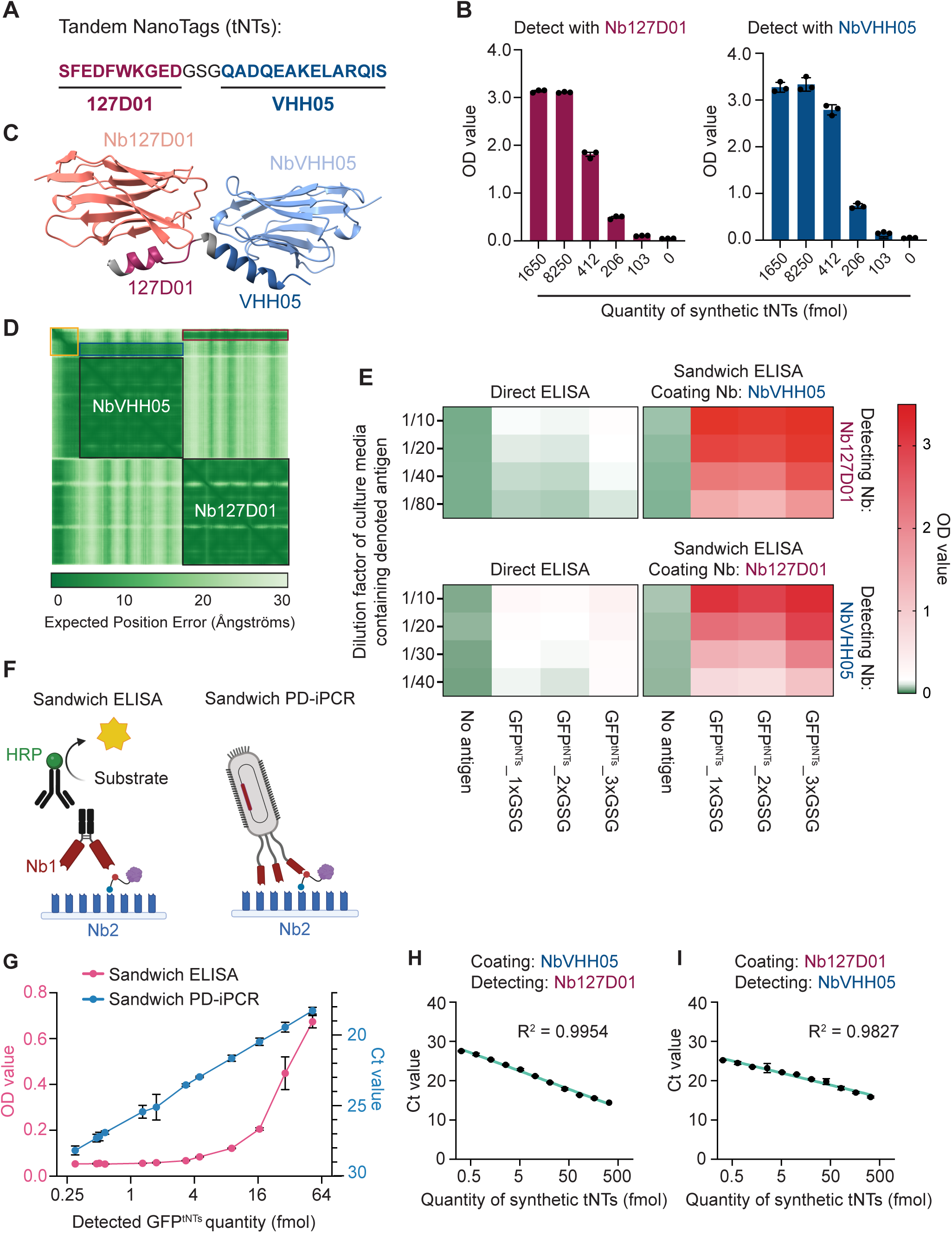
Establishment of phage display-mediated immuno-PCR with tandem NanoTags. (A) Amino acid sequence of tandem NanoTags. (B) Direct ELISA detecting tandem NanoTags with Nb127D01 or NbVHH05. N = 3. (C) AlphaFold3 model showing ternary protein complex composed of tandem NanoTags, NbVHH05, and Nb127D01. (D) Expected position error of the ternary protein complex in (*C*). Yellow, blue, and red squares indicate tandem NanoTags (GSG-127D01-GSG-VHH05), interacting area between tandem NanoTags and NbVHH05, and interacting area between tandem NanoTags and Nb127D01, respectively. (E) Heatmaps showing OD values from direct and sandwich ELISA detecting diluted S2R+ conditioned medium containing GFP^tNTs^_1×GSG, GFP^tNTs^_2×GSG, or GFP^tNTs^_3×GSG. N = 3. (F) Schematics of sandwich ELISA and sandwich PD-iPCR. (G) Sensitivity comparison between sandwich ELISA and sandwich PD-iPCR detecting GFP^tNTs^_1×GSG. error bars, SEM; N = 3. (H and I) Ct values obtained from sandwich PD-iPCR using femtomole quantities of synthetic tandem NanoTags. error bars, SEM; N = 2 or 3.

To further enhance detection sensitivity, we adapted sandwich ELISA to PD-iPCR format (Figure 4F). While conventional sandwich ELISA could not detect 0.3-4 fmol GFP^tNTs^, sandwich PD-iPCR retained linear quantification within this range and achieved superior detection sensitivity (Figure 4G). Moreover, interchanging the nanobody pairs in sandwich PD-iPCR yielded similar coefficients of determination (R²) (Figures 4H and 4I), demonstrating robustness and reproducibility of this assay. Collectively, these results establish a highly sensitive sandwich PD-iPCR platform capable of detecting tNTs-tagged antigens at femtomole levels, highlighting its potential for quantifying secreted proteins present at extremely low concentrations in nanoliter-scale hemolymph samples.

To apply this method for circulating ImpL2 detection in *Drosophila* hemolymph, we generated flies carrying a scarless CRISPR/Cas9-mediated knock-in of tNTs at the C-terminus of the endogenous *ImpL2* locus (*ImpL2^tNTs^*) (Supplementary Figures 3A-C). Because ImpL2 contains an N-terminal signal peptide, we inserted tNTs at the C-terminus to avoid interfering with secretion. This strategy also ensured tagging of all annotated ImpL2 isoforms.

Since generated *ImpL2^tNTs^* flies showed homozygous lethality, we questioned whether the tNTs disrupted ImpL2 function. Given that ImpL2 deficiency promotes systemic insulin signaling and consequently increases body weight (10), we analyzed fly body weight. As expected, flies homozygous or heterozygous for the null allele *ImpL2^Def20^* showed increased body weights in both sexes (Supplementary Figures 3D and 3E). However, *ImpL2^Def20/tNTs^* flies were viable and exhibited significantly reduced body weights compared to *ImpL2^Def20/Def20^* flies. The body weights of *ImpL2^Def20/tNTs^* flies were comparable to those of *ImpL2^Def20/+^*flies, and *ImpL2^tNTs/+^*flies showed similar body weights to control flies (Supplementary Figures 3D and 3E). These data suggest that ImpL2^tNTs^ protein is functional and the lethality of *ImpL2^tNTs/tNTs^* was not due to loss of ImpL2 function.

We then quantified circulating endogenous ImpL2^tNTs^ protein in the hemolymph of *ImpL2^tNTs/+^* flies using sandwich PD-iPCR. The assay detected 0.87 fmol and 1.37 fmol from 25 µg total hemolymph proteins from male and female flies, respectively, which corresponds to 1.04 nM and 1.42 nM ImpL2^tNTs^ protein (Supplementary Figures 3F and 3G; refer to Materials and Methods section for detailed molarity calculation). These data demonstrated that this approach enables sensitive and quantitative measurement of endogenous ImpL2 protein *in vivo*.

### Quantification of ImpL2 under various physiological conditions using sandwich PD-iPCR

ImpL2 protein has been reported to increase under starvation conditions, where it acts to conserve circulating nutrients by suppressing Dilp2 signaling (10, 15). To detect this increase, we performed sandwich PD-iPCR using hemolymph from fed and starved *ImpL2^tNTs/+^* flies and found that starved flies had 3.2-3.9 nM circulating ImpL2^tNTs^ protein, representing a 2.7-3.7-fold increase compared to fed flies (Figure 5A). This increase was also confirmed by Western blot analysis (Figure 5B), though the sensitivity was notably lower than that of sandwich PD-iPCR, emphasizing the superior detection performance of PD-iPCR.

**Figure 5.**
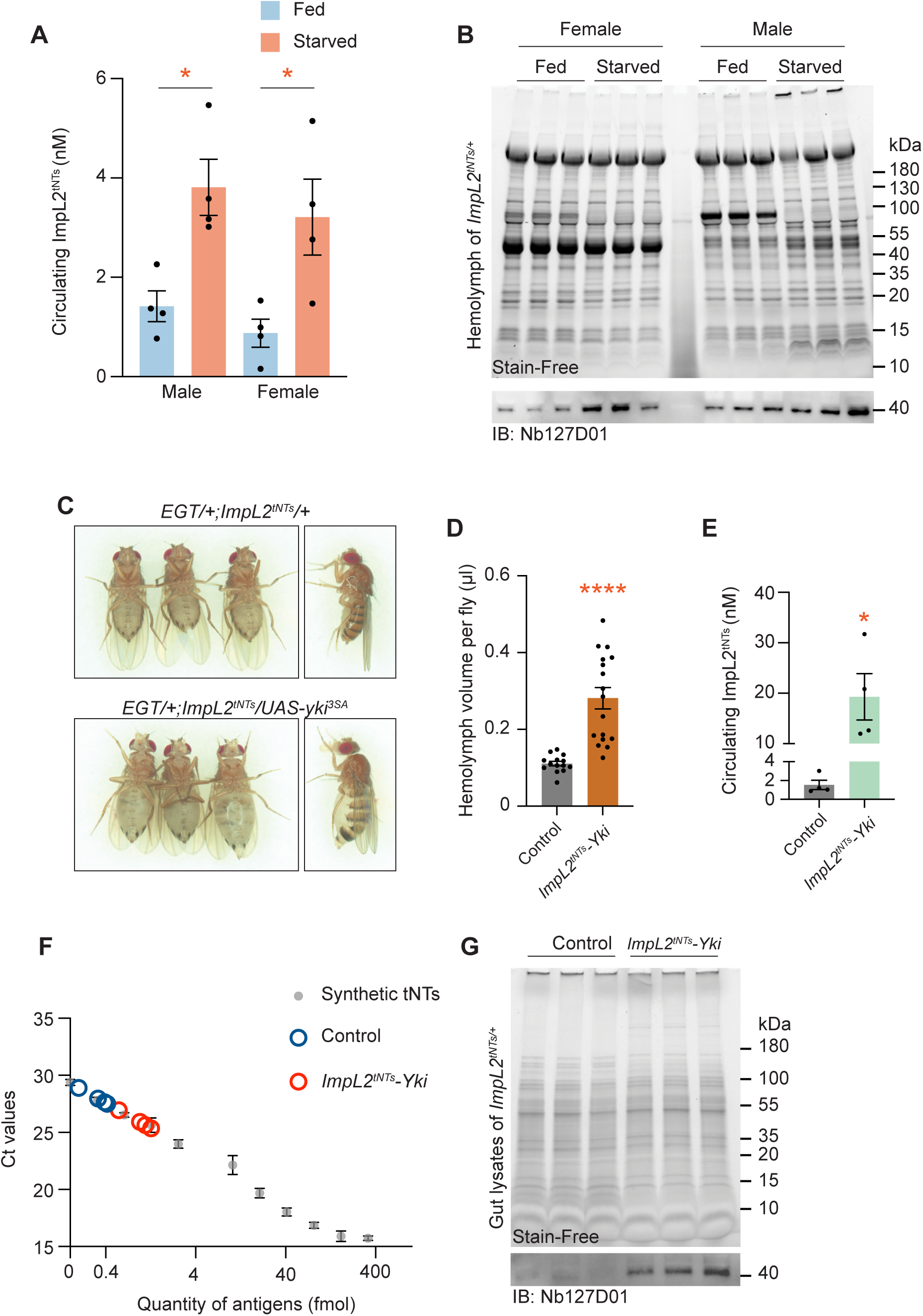
Detection of ImpL2^tNTs^ with sandwich PD-iPCR *in vivo*. (A) Molarity of circulating ImpL2^tNTs^ measured by sandwich PD-iPCR in the hemolymph of fed and starved *ImpL2^tNTs^*^/+^ flies. \**P* < 0.05 by Student’s t test; error bars, SEM; a single replicate = 50 µg proteins from pooled fly hemolymph; N = 4. (B) Western blotting showing ImpL2^tNTs^ proteins in the hemolymph of fed and starved *ImpL2^tNTs^*^/+^ flies. (C) Representative images showing bloating phenotype in *EGT/+; ImpL2^tNTs^/+* and *EGT/+; ImpL2^tNTs^/UAS-yki^3SA^* flies. (D) Hemolymph volume of control (*EGT/+; ImpL2^tNTs^/+*) and *ImpL2^tNTs^-Yki* (*EGT/+; ImpL2^tNTs^/UAS-yki^3SA^*) flies. \*\*\*\**P* < 1.0 × 10^−4^ by Student’s t test; error bars, SEM; control = 14 replicates and *ImpL2^tNTs^-Yki* = 17 replicates. (E) Molarity of circulating ImpL2^tNTs^ measured by sandwich PD-iPCR in hemolymph of control (*EGT/+; ImpL2^tNTs^/+*) and *ImpL2^tNTs^-Yki* (*EGT/+; ImpL2^tNTs^/UAS-yki^3SA^*) flies. \**P* < 0.05 by Student’s t test; error bars, SEM; a single replicate = 50 µg proteins from pooled female fly hemolymph; N = 4. (F) Sandwich PD-iPCR detecting ImpL2^tNTs^ in gut lysates of control (*EGT/+; ImpL2^tNTs^/+*) and *ImpL2^tNTs^-Yki* (*EGT/+; ImpL2^tNTs^/UAS-yki^3SA^*) flies. Standard curve was created from synthetic tNTs. \**P* < 0.05 by Student’s t test; error bars, SEM; a single replicate = 10 µg proteins from female gut lysates; N = 4. (G) Western blot showing ImpL2^tNTs^ in gut lysates of control (*EGT/+; ImpL2^tNTs^/+*) and *ImpL2^tNTs^-Yki* (*EGT/+; ImpL2^tNTs^/UAS-yki^3SA^*) flies.

We next applied the tNTs-based sandwich PD-iPCR to quantify ImpL2 levels in flies with *yki^act^*-driven gut tumors. Tumors were induced for 8 days in the gut of *EGT > yki^act^* flies carrying a single *ImpL2^tNTs^* allele (*EGT/+; ImpL2^tNTs^/UAS-yki^3SA^*, hereafter referred to as *ImpL2^tNTs^-Yki* flies). These flies exhibited a bloated phenotype and significantly increased hemolymph volume (Figures 5C and 5D), consistent with the previous *Yki^act^* tumor model (21). Using sandwich PD-iPCR, we measured 19.3 nM of ImpL2^tNTs^ protein in the hemolymph of *ImpL2^tNTs^-Yki* flies—a 12.5-fold increase compared to the controls (Figure 5E). When accounting for the 2.54-fold increase in hemolymph volume (Figure 5D), this translates to an overall 31.75-fold increase in total circulating ImpL2^tNTs^ in *ImpL2^tNTs^-Yki* flies.

Previously, *yki^act^* gut tumors have been suspected as the major source of increased ImpL2 based on their elevated *ImpL2* mRNA levels (21). However, protein-level confirmation has been lacking. We therefore performed sandwich PD-iPCR using 10 µg of total proteins from dissected gut lysates of *ImpL2^tNTs^-Yki* and control flies to detect ImpL2^tNTs^ proteins. Gut lysates from *ImpL2^tNTs^-Yki* flies exhibited noticeably lower Ct values compared to control gut lysates, whose Ct values were near the limit of detection of sandwich PD-iPCR, indicating higher ImpL2^tNTs^ protein levels in the gut of *ImpL2^tNTs^-Yki* flies compared to control flies (Figure 5F). The minimal detection of ImpL2^tNTs^ protein in control gut lysates is consistent with previous single-nucleus RNA sequencing data (24). This increase in ImpL2^tNTs^ protein level in gut of *ImpL2^tNTs^-Yki* flies was further confirmed by Western blotting (Figure 5G). Together, these results demonstrate that tNTs-based sandwich PD-iPCR enables sensitive, quantitative detection of endogenous ImpL2 protein at nanomolar level *in vivo* under various physiological and pathological conditions, including starvation and tumorigenesis.

## Discussion

In this study, we established PD-iPCR as a novel and highly sensitive method for detecting low-abundance *Drosophila* secreted proteins using two strategies: 1) the generation of high affinity nanobodies through screening and affinity maturation, and 2) tagging of target secreted protein with tNTs that are specifically recognized by their corresponding nanobodies. Using ImpL2 as an example, we demonstrated that PD-iPCR enables reliable detection and quantification of secreted ImpL2 protein in the hemolymph under various physiological conditions, including starvation and tumorigenesis. Notably, tNTs-based sandwich iPCR achieved picomolar-level detection sensitivity, highlighting its potential for quantifying secreted proteins that exist at extremely low concentrations within nanoliter volumes of hemolymph. This approach can be readily leveraged to study secreted protein dynamics *in vivo* with high sensitivity and specificity.

Through nanobody screening and affinity maturation, we identified two high-affinity nanobodies that specifically recognize ImpL2 (Figure 2), yielding valuable reagents for ImpL2 studies. Notably, during the affinity maturation of NbImpL2-2B, an asparagine-to-lysine (N-to-K) substitution in CDR2 enhanced binding affinity. However, this position was limited to asparagine (N) and tyrosine (Y) residues in the original library design (16), suggesting that the library’s diversity may have constrained the identification of higher-affinity binders. Expanding diversity of the library could increase the likelihood of discovering nanobodies with further improved affinity.

Despite the successful identification of nanobodies, the screening process remains technically challenging, as it involves purification of recombinant mature secreted proteins followed by multiple rounds of selection and affinity maturation. Furthermore, the nanobodies with an improved binding affinity were unable to detect endogenous ImpL2 in *yki^act^*tumor flies (Supplementary Figure 2). In contrast, tNTs-based PD-iPCR achieved picomolar detection sensitivity for the tagged proteins (Figs. 4H and 4I) and enabled quantification of endogenous ImpL2^tNTs^ under starvation and tumorigenic conditions (Figure 5). The present study focuses on ImpL2 as a proof-of-concept, and smaller peptide hormones, such as Akh, AstC, tachykinins, or NPF, may be more challenging to tag without perturbing their native function requiring empirical functional validation after tagging. Although the CRISPR-based tagging strategy requires validation to ensure that tagging does not disrupt native protein function, endogenous tagging is generally well-tolerated. For example, 77% of *Drosophila* lines expressing intronic GFP-tagged essential genes retained normal viability (25). Furthermore, numerous GFP-tagged proteins exhibited precise localization across tissues and subcellular compartments, indicating preserved protein activity (25). Because tNTs is a compact epitope consisting of only 27 amino acids, we anticipate that this approach will serve as a minimally invasive and highly sensitive tagging strategy for studying secreted proteins in *Drosophila*.

Moreover, PD-iPCR offers the capability to perform multiplex antigen detection using minimal sample volumes. With the recent development of high affinity nanobodies targeting conventional epitope tags such as HA, FLAG, ALFA, and V5 (26–29), this platform could be readily adapted to quantify multiple tagged proteins simultaneously from same fly hemolymph sample. For instance, a transgenic fly line expressing double-tagged Dilp2 (*Ilp2^HF^*) has been used for quantifying Dilp2 levels (6). Given that ImpL2 functions as a secreted antagonist of Dilp2 signaling, simultaneous measurement of ImpL2 and Dilp2 across diverse physiological contexts, such as varying dietary states or metabolic disease conditions, would provide valuable insights into their dynamic interplay. Furthermore, with numerous GFP-tagged *Drosophila* lines (25) and a collection of GFP nanobodies (30), the PD-iPCR platform is well positioned to expand its capacity toward multiplexed protein quantification *in vivo*. We expect that this platform would broaden our understanding on dynamic regulation of secreted proteins and hormone in *Drosophila*.

## Materials and Methods

### Cell culture and transfection

S2R+ cells were cultured in Schneider’s media (Thermo, 21720024) supplemented with 10 % FBS and 50 U/ml Penicillin-Streptomycin (Thermo, 15070063) at 25 °C. S2R+ cells were transfected with plasmids and dsRNA using Effectene (Qiagen, 301425) according to the manufacturer’s protocol. In brief, 3×10^6^ cells were plated in a well of a six-well plate prior to transfection. Either 400 ng plasmid DNA or 20 µg dsRNA was diluted in Buffer EC to 100 μL and mixed with 3.2 μL enhancer by vortexing. Then, 10 μL Effectene reagent was added to the mixture and vortexed again. After a 15-minute incubation at room temperature, the mixture was added dropwise to the *Drosophila* cells. Conditioned media were collected three days after transfection.

### Antibodies

The endogenous ImpL2 antibody used for immunoblotting was generously provided by Linda Partridge’s lab (8). Other antibodies used in this study include anti-M13 major coat protein antibody (Santa Cruz Biotechnology, sc-53004 HRP and sc-53004 AF647), anti-alpaca VHH domain antibody (Jackson ImmunoResearch, 128-605-230 and 128-035-230) and anti-Human IgG Fc antibody (Thermo Fisher Scientific, A18829).

### dsRNA synthesis

dsRNAs were designed by the *Drosophila* RNAi Screening Center and synthesized as previous described (31). dsRNA targeting GFP sequence was amplified from GFP plasmid using primers: Forward 5’-TAATACGACTCACTATAGGGAAGTTCATCTGCACCACCGG-3’ and Reverse 5’-TAATACGACTCACTATAGGGTCTTTGCTCAGGGCGGACTG-3’. The ImpL2-ds1 (DRSC28691) was amplified from *Drosophila* genomic DNA using primers: Forward 5’-TAATACGACTCACTATAGGGGGTGTAGATGATTCGCGGTT-3’ and Reverse 5’-T TAATACGACTCACTATAGGGTTACATGTGTGCGCCTTAGC-3’. The ImpL2-ds2 (DRSC08666) was amplified from *Drosophila* genomic DNA using primers: Forward 5’-TAATACGACTCACTATAGGGGATCTCCTTGTTCTCGTTATTC-3’ and Reverse 5’-T TAATACGACTCACTATAGGGCCTTCGAAGCCGACTGG-3’. dsRNAs were synthesized using MEGAscript T7 Transcription Kit (Invitrogen) and purified using RNeasy Mini Kit (QIAGEN).

### Fly husbandry

Flies were maintained on standard cornmeal food and raised at 25 °C with 12 h:12 h light: dark cycles. For the *ImpL2*-overexpression experiment, *Tub-Gal80^TS^/UAS-empty;mef2-Gal4/+* and *Tub-Gal80^TS^/+;mef2-Gal4/UAS-ImpL2* were induced at 29 ℃ for two days to induce *ImpL2* expression. For the gut tumor experiment, *EGT/+*, *EGT/+;UAS-yki^3SA^/+, EGT/+;ImpL2^tNTs^/+*, and *EGT/+;ImpL2^tNTs^/UAS-yki^3SA^* flies were raised at 18 °C until 1 day after eclosion and transferred to 29 °C to induce expression of *yki^3SA^*. During 29 °C culture, flies were transferred to fresh food vial every 2 days. Starved flies were induced by cultivating the flies in empty vials containing cotton ball soaked in PBS for 2 days. The following fly stocks were used in this study: *UAS-Yki^3SA^* (32), *UAS-ImpL2* (10), *UAS-empty* (BDSC# 36304) and *ImpL2^Def20^* (gift from Young Kwon). *yw*, *Tub-Gal80^TS^;mef2-Gal4*, and *EGT* (*esg-GAL4, UAS-GFP, tub-GAL80^TS^*) are from the Perrimon lab stock collection.

### ImpL2-hIgG purification from conditioned medium using stable S2 cells

ImpL2 was purified from conditioned medium following protocol modified from our previous report (16). The protein sequence of secreted ImpL2 is as follows: RAVDLVDDSNDVDNSIEAEEEKPRNRAFEADWLKFTKTPPTKLQQADGATIEIVCEMMGSQVPSIQWVVGHLPRSELDDLDSNQVAEEAPSAIVRVRSSHIIDHVLSEARTYTCVGRTGSKTIYASTVVHPPRSSRLTPEKTYPGAQKPRIIYTEKTHLDLMGSNIQLPCRVHARPRAEITWLNNENKEIVQGHRHRVLANGDLLISEIKWEDMGNYKCIARNVVGKDTADTFVYPVLN EEDQL. Briefly, *Drosophila* S2 cells cultured in serum-free ESF921 medium (Expression Systems, 96-001-01) were transfected with a plasmid encoding ImpL2-hIgG driven by metallothionein promoter and selected with puromycin to establish a stable cell line. Then, the cells are adapted to suspension culture to enhance protein yield. ImpL2-hIgG expression was induced by adding 500 µM CuSO_4_, and after 4-5 days culture, the conditioned medium (CM) was collected by centrifugation at 1000 ×g, for 10 min and filtrated with 0.22 µm membrane.

To purify ImpL2-hIgG from CM, 1 mL of Protein A resin (GenScript, L00210) was equilibrated with 25 mL TBS in a gravity column. Next, the column was loaded with 50-250 mL of CM, washed three times with TBS, and eluted with 5 mL of 0.1 M glycine (pH 3.5). The eluates were immediately neutralized with 0.5 mL of Tris-HCl (pH 8.5) and pooled with an additional 2 mL TBS rinse of the resin. The elutes were concentrated using an Amicon Ultra-15 filter (50 kDa cutoff; Millipore Sigma) at 4150 ×g for 30 min at 4 °C, with buffer exchanges into TBS for 3 times. Protein concentration was determined at A280 using a NanoDrop spectrophotometer or Bicinchoninic Acid (BCA) Assay (Thermo, 23227). Purified mCherry-hIgG was also prepared through same procedure.

### ImpL2 nanobody screening using phage-displayed nanobody library

Purified ImpL2-hIgG proteins were diluted in PBS and coated onto Nunc MaxiSorp 96-well plates (Thermo Fisher Scientific, 442404) overnight at 4 °C (Coated ImpL2-hIgG quantity: 1 µg per well in round 1, 0.5 µg in round 2, 0.25 µg in round 3). Plates were then blocked for 1-2 h at room temperature (RT) with 5 % MPBST (5 % skim milk in PBS with 0.05 % Tween-20). For negative selection, mCherry-hIgG-coated wells (1 µg per well) were prepared in parallel under identical conditions. The phage displayed nanobody libraries were diluted with 5 % MPBST and first incubated with mCherry-hIgG plates (60 µL per well) at room temperature for 1 hour to remove nonspecific binders. The unbound fraction was then transferred to ImpL2-hIgG-coated wells. Residual unbound fractions in the negative selection plates were collected with additional 90 µL 5 % MPBST rinses and pooled to 60 µl phages in positive selection plates (final 150 µL per well) and incubated for 1 hour at room temperature. Plates were washed with TBST (1×TBS with 0.1 % Tween-20). The TBST washing strength was gradually increased in each round (10×, 20×, and 30× in rounds 1-3, respectively). The bound phages were eluted by infecting *E. coli* OmniMAX (Invitrogen, C854003; OD600 at 0.6) for 30 min at RT with gentle shaking, followed by helper phage infection (MOI at 20) for 1 hour at RT. The infected bacteria were cultured overnight in 2×YT medium supplemented with ampicillin (100 µg mL^-1^) and kanamycin (50 µg mL^-1^) at 37 °C with shaking at 220 rpm. The following day, phages were precipitated from the culture supernatant using PEG/NaCl solution (20 % PEG8000, 2.5 M NaCl) and pelleted by centrifugation at 12,000 ×g for 30 min at 4 °C. The resulting pellet was resuspended in 2 mL PBS. 1 mL phage solution was used for the next round screening, and the remaining solution was kept for polyclonal ELISA.

### ELISA with polyclonal phages

ImpL2-hIgG and mCherry-hIgG proteins were serially diluted, coated on MaxiSorp plates, and blocked with 5 % MPBST for 1 hour. Polyclonal phages from the first, second, and third selection rounds were incubated with the antigen-coated plates for 1 hour at room temperature, washed with TBST, and probed with anti-M13-HRP (Santa Cruz, sc-53004) for 1 hour at room temperature. After additional TBST washes, 100 μL TMB substrate (Thermo Fisher, 34028) was added, and the reaction was stopped with 50 μL 0.4 M H_2_SO_4_. Absorbance at 450 nm was measured with plate reader.

### ELISA with monoclonal phages

96 bacteria colonies from the third selection round were picked into deep-well plates containing 100 µL 2×YT with carbenicillin and grown at 37 °C for 6-8 hours until OD600 reaches to 0.6. Cultures were infected with helper phages for 30 min at room temperature (RT), supplemented with 1 mL 2×YT containing carbenicillin and kanamycin, and cultured overnight at 37 °C, 220 rpm.

Monoclonal phage supernatants were harvested and used as primary antibodies in ELISA. MaxiSorp plates were coated with ImpL2-hIgG or mCherry-hIgG (0.25 µg/well) and blocked as described above. Phage supernatants diluted with same volume of blocking buffer were added to the plates and incubated for 1 hour at RT. After incubation, the plates were washed 10 times with TBST and incubated with anti-M13 major coat protein-HRP antibody for 1 hour. After three times washing with TBST, 100 µL TMB substrate was applied, and the reaction was stopped with 50 µL 0.4 M H_2_SO_4_. Absorbance at 450 nm was measured with plate reader.

### Purification of recombinant nanobody

Nanobodies were expressed in bacteria and secreted into bacteria periplasm using pET-26b-Nb-GGA plasmid as previous described (3, 16). The periplasm fraction was extracted using B-PER II reagent (Thermo Fisher Scientific, 78260) and loaded onto a gravity column containing 1 mL Ni²⁺-NTA resin (Cytiva, 17531801) pre-equilibrated in TBS with 10 mM imidazole. The column was washed three times with 20 mL TBS containing 20 mM imidazole, and the bound nanobodies were eluted stepwise in 1 mL TBS supplemented with 100, 250, and 500 mM imidazole. The elutes were analyzed by SDS-PAGE, and those containing the nanobodies were pooled, dialyzed, and stored at -80 °C.

### Phage purification

One Shot® OmniMAX™ 2 T1 Phage-Resistant Cells (Thermo Fisher Scientific, C854003) transformed with phagemids expressing nanobody-M13 pIII fusion proteins were cultured in Carbenicillin containing 2×YT media (10 g/L yeast extract, 16 g/L Tryptone, and 5 g/L NaCl) until OD600 reaches to 0.6. 500 µL cultures were infected with 50 µL M13K07 helper (Thermo Fisher Scientific, 18311019) for 1 hour at 37 °C. Then, the 500 µL culture was mixed with 500 mL 2×YT medium (+ Carb and Kan) and cultured overnight at 37 °C. After spinning down the bacterial culture, phage in the culture supernatant was precipitated following addition of a quarter volume of PEG/NaCl solution (20 % PEG-8000 (Thermo Fisher Scientific, BP233-1) and 2.5 M NaCl) and subsequent ice incubation for 20 min. Phage pellets were acquired after 12,000 ×g centrifugation for 30 min and resuspended in PBS.

### Immunostaining of ImpL2-GPI cells using nanobody-displaying phages or purified nanobodies

S2R+ cells were transfected with plasmids co-expressing ImpL2-GPI and GFP. Two days after transfection, live cells were incubated in blocking buffer (5 % skim milk in PBS) for 30 min at 4 °C without fixation and permeabilization. Next, they were incubated with nanobody-displaying phages or purified nanobodies for 1 hour at 4 °C. After incubation, cells were washed three times with PBS and stained for 30 min at 4 °C using either anti-M13 major coat protein-Alexa Fluor 647 (Santa Cruz Biotechnology, sc-53004 AF647) or anti-VHH-Alexa Fluor 647 (Jackson ImmunoResearch, 128-605-230). Following three additional PBS washes, samples were imaged on IN Cell high-throughput confocal microscope (MicRoN Core, Harvard Medical School).

### Nanobody affinity maturation by random mutagenesis

The random mutagenesis was performed by PCR using the GeneMorph II Random Mutagenesis Kit (Agilent, 200550). The CDR and framework regions were amplified from each nanobody phagemid using the following primers: Forward 5’-TGAGCTGCGCGGCGAGCGGC-3’ and Reverse 5’-CACCTGGGTGCCCTGGCCCC-3’. The PCR products were assembled into the phagemid vector pLibF using the NEBuilder HiFi DNA Assembly Master Mix (NEB, E2621L). The resulting constructs were electroporated into TG1 cells to generate mutant libraries for each nanobody. The transformed bacteria were cultured to an OD600 of 0.6, infected with helper phage at an MOI of 20, and incubated overnight in 2×YT medium supplemented with ampicillin (100 µg/mL) and kanamycin (50 µg/mL) at 37 °C with shaking at 220 rpm. The next day, phages were collected from the culture supernatant by PEG/NaCl precipitation (20 % PEG8000, 2.5 M NaCl) and pelleted by centrifugation at 12,000 ×g for 30 min at 4 °C. The pellet was resuspended in PBS and used for subsequent round of screening using mCherry-hIgG as negative selection and ImpL2-hIgG as positive selection. The bound phages were recovered by infecting *E. coli* OmniMAX. After overnight culture, the phagemids were purified using the Spin Miniprep Kit (Qiagen, 27106) and used as PCR templates for a second round of mutagenesis to construct the second mutant libraries for each nanobody. After each selection cycle with these libraries, monoclonal phages were isolated and analyzed by ELISA.

### Generation of ImpL2^tNTs^ fly

To generate *ImpL2^tNTs^*, we followed a modified version of the protocol described before (33) Homology donor intermediate vectors were ordered from Genewiz in the pUC57 Kan_gw_OK2 vector backbone, containing a gRNA targeting the 3’ end of *ImpL2*, a linker and 127D01-tag, 200 bp short homology arms flanking the genomic cut site, and a BsaI cloning site (Supplementary Figure 3A) to produce the ImpL2-pUC57 plasmid. A silent mutation was introduced into the non-coding sequence in the homology arm PAM to avoid cleavage of the plasmid. Insert DNA containing linker, VHH05-tag, and PBac scarless cassette was PCR amplified from pScarlessHD-C-3xVHH05-DsRed plasmid (Addgene, 171580) using the following primers:

Fwd 5’-ATCTGGTCTCAGGTTCGGGACAGGCCGATCAGG-3’

Rev 5’-ATCGGTCTCATTAACCCTAGAAAGATAGTCTG-3’

Both the ImpL2-pUC57 plasmid and PCR product were digested with BsaI and ligated with 2.5 µL 10× T4 DNA ligase buffer (NEB, B0202S) and 0.5 µL T4 DNA ligase (NEB, M0202S) to produce the finished construct, ImpL2-tandem-nanotag (Supplementary Figure 3B). The construct was injected (250 ng/µl) into 300 *yw; attP40, nos-Cas9* embryos as described previously (34) Resulting G0 males and females were crossed individually to *yw; Dr e/TM3, Sb* flies for 3×P3-RFP screening. Positive lines were balanced, and stocks were established. To produce the finished scarless C-terminal tag, the 3×P3-RFP marker cassette was removed by crossing to a fly line expressing the piggy Bac transposase (BDSC, 8283). To verify the insertion, PCR primers that flank the integration site were used in combination with primers that bind within the inserted cassette in both orientations. 500-800 nt amplicons were amplified from genomic DNA from individual insertion lines through single fly PCR using GoTaq green master mix (Promega, M7122). Following balancing of *ImpL2^tNTs^* with *TM6B*, no progeny homozygous for the *ImpL2^tNTs^*allele were recovered.

### Hemolymph collection

30-50 flies were pierced with a tungsten needle and collected into pierced 0.5 mL Protein LoBind® Tubes (Eppendorf, 022431064). The 0.5 mL Eppendorf tube containing the pierced flies was placed in 1.5 mL Protein LoBind® Tube (Eppendorf, 022431081) and was centrifuged at 9,000 ×g for 5 min at 4 °C following previously reported protocols (6, 35). The collected fly hemolymph in the 1.5 mL Eppendorf tube was mixed with 1×PBS as a diluent (36–38) containing 1×cOmplete™, EDTA-free Protease Inhibitor Cocktail (Roche, 04693132001) and 1×PhosStop (Roche, 04906845001), and was additionally centrifuged to remove hemocytes and debris. The volume of collected hemolymph was measured with micropipette before adding 1×PBS.

### Direct or sandwich ELISA

For sandwich ELISA, Maxisorp 96 well ELISA plate (Thermo Fisher Scientific, 442404) was coated with NbVHH05 or Nb127D01 nanobodies (22) diluted in 1×PBS (50 ng per well) overnight at 4 °C. Then, the plate was washed with 200 µL TBST three times and subsequently blocked with 200 µL of 5 % BTBST blocking buffer (TBST containing 5 % bovine serum albumin) for 1 hour at 25 °C. After aspirating the 5 % BTBST blocking buffer, 100 µL of antigen diluted in 5 % BTBST was treated for 1 hour at RT followed by three times of washing with TBST. Then, the ELISA plate was incubated for 1 hour with 100 µL of Nb127D01-human IgG or NbVHH05-human IgG (1:100 dilution of S2 cell conditioned media containing the nanobodies in 5 % BTBST) to detect the antigens tagged with tandem NanoTags. After three times of washing with TBST, 100 µL of Goat anti-human IgG-HRP (1:5000 diluted in 5 % BTBST, Thermo Fisher Scientific, A18829) was treated for 1 hour at 25 °C. After washing with TBST, 100 µL of TMB ELISA Substrate Solutions (Thermo Fisher Scientific, PI34029) was treated for 15 min, and the absorbance was measured with a plate reader at 650 nm. Direct ELISA was performed by coating the Maxisorp ELISA plate with the antigens tagged with tandem NanoTags for overnight at 4 °C. The next day, the plate was washed with 200 µL TBST three times and subsequently blocked with 200 µL of 5 % BTBST blocking buffer for 1 hour at 25 °C. After blocking, the plate was incubated with nanobodies diluted in 5 % BTBST and processed as described in the sandwich ELISA protocol.

### Phage display-mediated sandwich immuno-PCR

Coat the Maxisorp-treated 96 well ELISA plate (Thermo Fisher Scientific, 442404) with 2 µg of purified capture nanobody (50 µL of 40 µg/mL nanobody diluted in 1×PBS per well) overnight at 4 °C. The ELISA plate was washed with TBST (0.1 % Tween-20 in 1×PBS) three times and incubated with 200 µL of 5 % MTBST blocking buffer for 1 hour at RT. After removing 5 % MTBST, 50 µL of antigen diluted in 5 % MTBST was treated for 1 hour at RT. The antigen was aspirated, and ELISA plate was washed with TBST three times. Then 50 µL of detection nanobody displaying phages diluted in 5 % MTBST was treated for 1 hour at RT. After the incubation with nanobody-displaying phages, the ELISA plate is washed with TBST ten times and subsequently washed with distilled water three times. Then, the phage bound to the ELISA plate was eluted with 50 µL of 0.2 M glycine (pH 2.5) for 20 min at RT and immediately neutralized by adding 5 µL of 1 M Tris (pH 8.5). To detect the phage DNA with quantitative-PCR (qPCR), forward primer 5’-GCGCTACAGTCTGACGCTAA-3’ and reverse primer 5’-ACCATTAGCAAGGCCGGAAA-3’ were used. The qPCR reaction was prepared with the mixture composed of iQ™ SYBR® Green Supermix (Bio-Rad, 171-8880), 0.25 µM forward primer, 0.25 µM reverse primer, and 4.5 µL phage eluate and analyzed with CFX96 Real-Time PCR Detection System (Bio-Rad). Ct value of each sample was averaged by two or three technical replicates. In every tNTs-based PD-iPCR experiment, duplicates or triplicates of serially diluted synthetic tNTs (SFEDFWKGEDGSGQADQEAKELARQIS, synthesized from GenScript) were included, and the molarity of ImpL2^tNTs^ was quantified based on the Ct values from the synthetic tNTs.

To quantify ImpL2^tNTs^ molar concentration, hemolymph was extracted from ∼100 adult flies carrying *ImpL2^tNTs/+^* transgene. Hemolymph was extracted by centrifugation, and the recovered volume was measured using micropipette. The hemolymph was then diluted into 10 or 20 µl of 1×PBS supplemented with 1×cOmplete™, EDTA-free Protease Inhibitor Cocktail (Roche, 04693132001) and 1×PhosStop (Roche, 04906845001). Total protein concentration of the diluted hemolymph sample was determined with BCA assay. For each biological replicate, 25 or 50 µg of total hemolymph protein was used as input for sandwich PD-iPCR. Based on the dilution factor and the amount of total protein loaded, the corresponding volume of original hemolymph loaded in each sandwich PD-iPCR was calculated. After performing sandwich PD-iPCR, Ct value obtained from of hemolymph samples was converted to absolute quantities of ImpL2^tNTs^ protein using a standard curve of synthetic tNTs. Finally, the molar concentration of ImpL2^tNTs^ protein in hemolymph was calculated by dividing the quantified amount of ImpL2^tNTs^ protein by the estimated volume of original hemolymph loaded into the reaction.

### Structure prediction with AlphaFold

For AlphaFold-Multimer predictions of anti-ImpL2 Nanobodies (NbImpL2-2B and NbImpL2-2B-M2-8D) complexed with ImpL2, we used LocalColabFold v.1.5.5 (39), which integrates AlphaFold-Multimer (AFM) v.2.3.1 (REF: https://www.biorxiv.org/content/10.1101/2021.10.04.463034). MSAs were generated using colabfold_search against the UniRef30 (version 2202) and mmseqs2/14-7e284 structural databases. Computations were performed on the Harvard O2 high-performance computing cluster. Each prediction generated five models with five recycling. Using the best predicted models from AlphaFold-Multimer, local interaction residues in contact-interface (Predicted Aligned Error ≤12 Å and Cβ–Cβ distance ≤ 8 Å) were identified via integrated Local Interaction Score analysis pipeline (40), available at https://github.com/flyark/AFM-LIS. The predicted structures were visualized in UCSF ChimeraX v1.9 (41).

Ternary structure composed of tNTs, NbVHH05, and Nb127D0 in Figure 4C was predicted with AlphaFold3 (https://alphafoldserver.com/). Input sequence is as follows:

tNTs: GSGSFEDFWKGEDGSGQADQEAKELARQIS

NbVHH05:

QVQLQESGGGLVQPGGSLRLSCAASGFVFENSAMAWYRQAPGKERELIAVIGTTFIKLAESVKGRFTISRDNAKSTVYLQMNNLKPEDTAVYYCSKSGAYWGQGTQVTVSS

Nb127D01:

EVQLVESGGGLVQAGESLRLSCAASGSTFDFKVMGWYRQPPGKQREGVAAIRLSGNMHYAESVKGRFAISKANAKNTVYLQMNSLRPEDTAVYYCKVNIRGQDYWGQGTQVTVSS

Western blot

Ten dissected fly guts were lysed in 45 µL of 1×Cell Lysis Buffer (Cell Signaling Technology, 9803) containing 1×cOmplete™, EDTA-free Protease Inhibitor Cocktail (Roche, 04693132001) and 1×PhosStop (Roche, 04906845001), and centrifuged at 14,000 ×g for 10 min, 4 °C. Protein concentration of the collected gut lysates or fly hemolymph was measured with BCA assay, and the same protein quantity was mixed with 4× Laemmli protein sample buffer (Bio-Rad, 1610747) and 2.5 % beta-mercaptoethanol followed by boiling at 95 °C for 5 min. Prepared protein samples were electrophoresed in 4-20 % Mini-PROTEAN® TGX Stain-Free™ Protein Gels (Bio-Rad, 4568093) and transferred to 0.2 µM nitrocellulose membrane (Bio-Rad, 1620112) with Trans-Blot® Turbo™ (Bio-Rad). The nitrocellulose membrane was blocked with TBST containing 5 % skim milk for 1 hour at RT. To detect the VHH05 or 127D01 epitopes, NbVHH05-human IgG or Nb127D01-human IgG (1:100, S2 cell culture media) and Goat anti-Human IgG-HRP (1:5000, Thermo Fisher Scientific, A18829) were used. The incubated antibodies were washed with TBST for 20 min, and signals were detected with enhanced chemiluminescence (ECL) (Thermo Fisher Scientific, 34095) and Bio-Rad ChemiDoc^TM^ MP imaging System.

## Acknowledgements and funding sources

We thank Young Kwon (University of Washington, USA) for providing the *ImpL2^Def20^* fly line, Afroditi Petsakou for providing the *ImpL2-OE* flies and Christians Villalta for performing injection. We are grateful to the Research Computing Group at Harvard Medical School for GPU access in the Harvard O2 high-performance computing cluster. This work is supported by NIH NIGMS P41 GM132087 (N.P.); the Cancer Grand Challenges partnership funded by Cancer Research UK (CGCATF-2021/100022), and the National Cancer Institute (1 OT2 CA278685-01). N.P. is an investigator of Howard Hughes Medical Institute.

This article is subject to HHMI’s Immediate Access to Research policy, which requires that this article be made publicly available as initial and revised preprints deposited on a designated preprint server under a CC BY 4.0 license.

## Author contributions

M.H., B.X., A.K., and N.P. conceived the study. M.H. established phage display-mediated iPCR with tandem NanoTag and conducted relevant experiments and data analysis. B.X. performed ImpL2 nanobody screenings and validation and conducted the experiments and data analysis for phage display-mediated iPCR with endogenous ImpL2 nanobodies. E.F. and J.Z. generated and established the *ImpL2^tNTs^*^/*Tm6B*^ fly line. A.K. and E.S. purified antigens and nanobodies. A.K. performed immunoprecipitation with ImpL2 nanobodies. M.H., T.M., and Y.L. conducted fly genetics and prepared fly samples. M.H., B.X., and N.P., wrote the manuscript.

---

**Supplementary figure 1.**
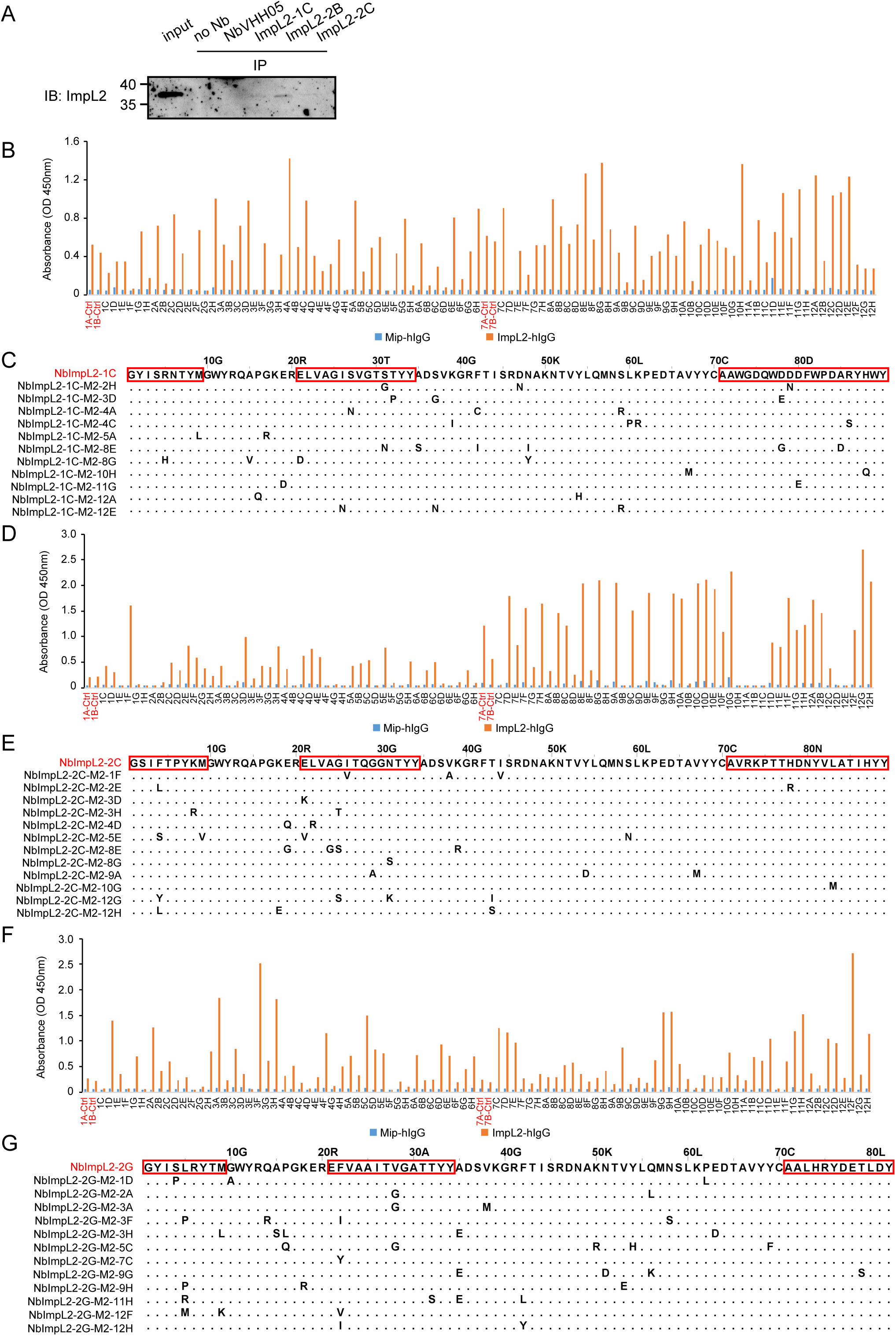
ImpL2 nanobody affinity maturation. (A) Immunoprecipitation of ImpL2 from S2R+ cell conditioned medium using ImpL2 nanobodies. NbVHH05 was used as a negative control. The ImpL2 was detected using endogenous ImpL2 antibody. (B) ELISA results of monoclonal phages isolated after two rounds of mutagenesis and selection of NbImpL2-1C. 96 individual clones were tested against the negative control protein (Mip-hIgG) and the target protein (ImpL2-hIgG). Clone 1A, 1B, 7A, 7B are phages displaying the original NbImpL2-1C nanobody. (C) Sequence alignment of NbImpL2-1C variants showing improved ELISA signals in (B). Mutations in the variants are indicated. The CDR regions are outlined with red rectangles. (D) ELISA results of monoclonal phages isolated after two rounds of mutagenesis and selection of NbImpL2-2C. 96 individual clones were tested against the negative control protein (Mip-hIgG) and the target protein (ImpL2-hIgG). Clone 1A, 1B, 7A, 7B are phages displaying the original NbImpL2-2C nanobody. (E) Sequence alignment of NbImpL2-2C variants showing improved ELISA signals in (D). Mutations in the variants are indicated. The CDR regions are outlined with red rectangles. (F) ELISA results of monoclonal phages isolated after two rounds of mutagenesis and selection of NbImpL2-2G. 96 individual clones were tested against the negative control protein (Mip-hIgG) and the target protein (ImpL2-hIgG). Clone 1A, 1B, 7A, 7B are phages displaying the original NbImpL2-2G nanobody. (G) Sequence alignment of NbImpL2-2G variants showing improved ELISA signals in (F). Mutations in the variants are indicated. The CDR regions are outlined with red rectangles.

**Supplementary figure 2.**
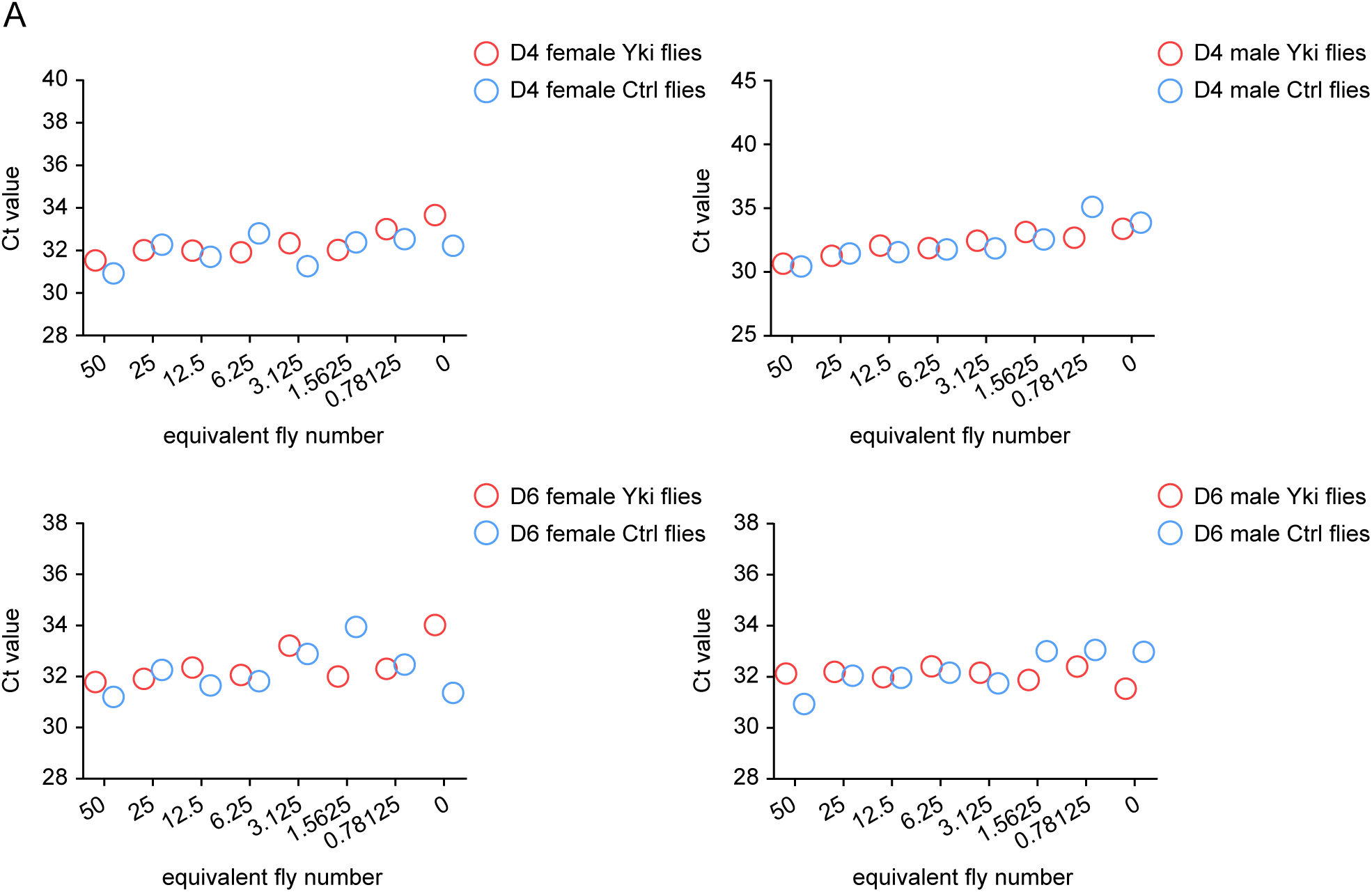
Quantification of ImpL2 in the hemolymph of flies bearing *Yki*-induced gut tumors. (A) Hemolymph was collected from Ctrl flies (*EGT/+;+/+*) and Yki flies (*EGT/+; UAS-yki^3SA^/+*) after four and six days of *yki^act^* induction. 2-fold serially diluted hemolymph samples were analyzed. Data represent one biological replicate.

**Supplementary figure 3.**
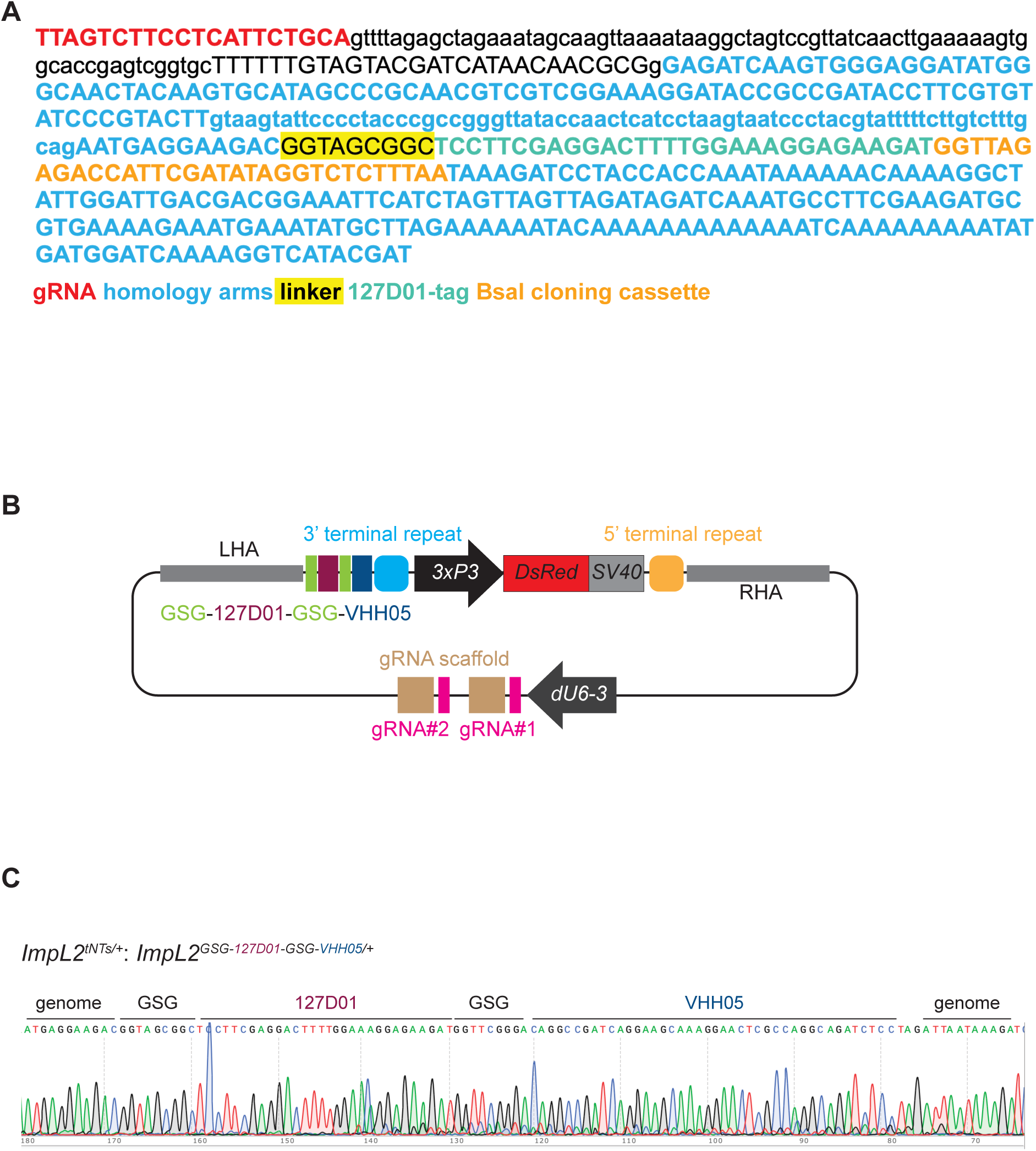
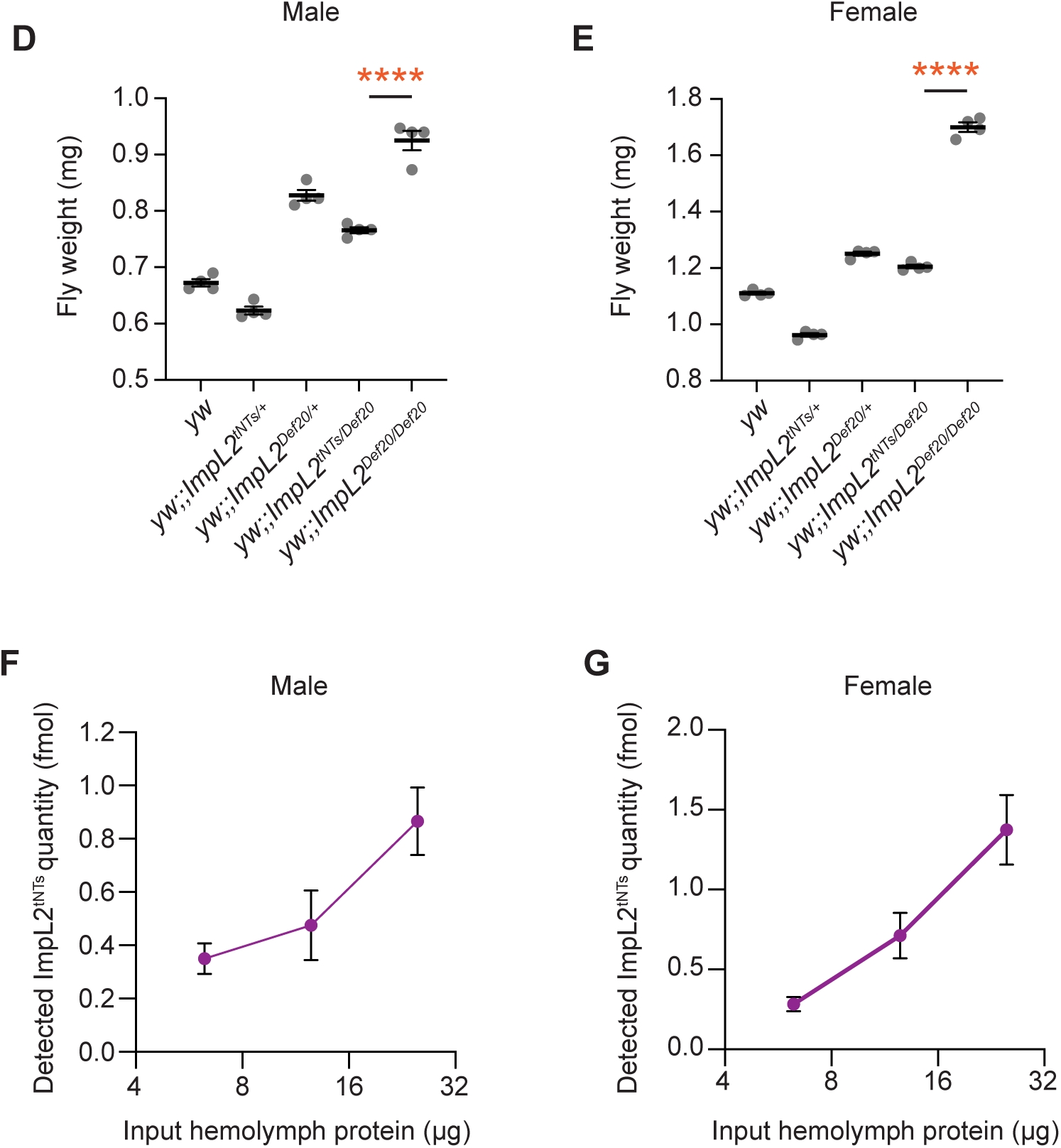
Generation of transgenic fly carrying knock-in of tandem NanoTags on *ImpL2* locus. (A) Sequence of the synthesized DNA cloned in pUC57 Kan_gw_OK2. (B) ImpL2-tandem-NanoTag plasmid map. (C) Sanger sequencing to verify the integration of tandem NanoTags in the C-terminal *ImpL2* locus. (D and E) Body weights of flies with the denoted genotype. \*\*\*\**P* < 1.0 × 10^−4^ by One-way ANOVA; error bars, SEM; N = 3. (F and G) ImpL2^tNTs^ quantity (fmol) from denoted input hemolymph proteins measured by sandwich PD-iPCR. X axis indicates the amount of total hemolymph proteins used. error bars, SEM; N = 3.

